# Laboratory Yeast Strains Rely On Oxidative Phosphorylation For Efficient ATP Production

**DOI:** 10.1101/2021.03.03.433835

**Authors:** Akshay Moharir, Lincoln Gay, Markus Babst

## Abstract

Even though it is a well-accepted fact that the energy metabolism of yeast is likely to impact all cellular activities, surprising little is known about the ATP homeostasis of particular yeast strains that are commonly used in cell biological studies. Therefore, we determined key parameters such as oxygen consumption and fermentation rates of the lab strain SEY6210. Our data indicated that even at high glucose concentrations, SEY6210 produces 30-50% of cellular ATP from oxidative phosphorylation. Loss of respiration, either by disrupting ATP synthase function or by growth in anaerobic conditions, was not fully compensated by fermentation and as a result affected energy intensive processes such as the maintenance of the plasma membrane proton gradient and the associated import of nutrients.

## Introduction

All cellular functions are directly or indirectly regulated by the cell’s energy metabolism. In particular, single-cell organisms such as yeast *Saccharomyces cerevisiae* have to be able to rapidly adjust cellular activities to changes in nutrient availability and other environmental conditions that impact energy levels. It is therefore surprising how little attention is given by cell biological studies to the energy status of the research subject. From our own experience, we know that even small changes in growth conditions of yeast can affect the outcome of the experiment (e.g., changing the diameter of the culture tube). These issues are particularly obvious when studying the trafficking of members of the Amino Acid Polyamine Organocation (APC) protein superfamily, which represent a major group of nutrient transporters that are responsible for the import of amino acids and other small molecules from the extracellular medium (reviewed in (Hundal and Taylor 2009)). Yeast expresses ~26 APC transporters, which use the strong proton gradient across the plasma membrane to import nutrients. This gradient is maintained by the P2-type H^+^-ATPase Pma1. With over 10^6^ molecules per cell, Pma1 is the most abundant protein in yeast and represents one of the main ATP consumers (Perlin, Harris et al. 1989, Cyert and Philpott 2013). It is therefore not surprising that Pma1 activity is tightly controlled by the metabolic state of the cell (Goossens, de La Fuente et al. 2000). As a consequence, APC transporters, which are major proton importers, have to be regulated parallel to Pma1 to prevent acidification of the cytoplasm and ensure that the proton gradient across the plasma membrane does not collapse. This rapid regulation of transporter activity is accomplished by regulating the rate of endocytosis and degradation of these proteins (Galan, Moreau et al. 1996, Seron, Blondel et al. 1999, Keener and Babst 2013, Ghaddar, Merhi et al. 2014, Gournas, Saliba et al. 2017).

In the attempt to understand the observed effects of stress conditions on APC transporter trafficking, we searched the literature for information about the energy metabolism of yeast strains commonly used for cell biological studies. However, we were unsuccessful and therefore we decided to measure the key parameters that determine the energy metabolism of lab strains.

## Results

The literature highlights the fact that at glucose concentrations commonly used in labs (2%), yeast relies mainly on fermentation for the synthesis of ATP (De Deken 1966, Johnston and Kim 2005). Oxidative phosphorylation is suppressed under high-glucose conditions and only is de-repressed when glucose is depleted (diauxic shift; (Galdieri, Mehrotra et al. 2010)). However, the findings of metabolic studies are often difficult to generalize, because of the use of different yeast strains, growth conditions and methods to determine changes in the metabolic state. Therefore, we decided to analyze the metabolic behavior of lab strains and growth conditions commonly used for cell biological studies.

### Oxygen consumption of lab strains

We determined the oxygen consumption of the wild-type strains SEY6210, BY4741 and W303 in rich medium containing 2% glucose (YPD medium). The cells were grown to mid-log phase and the oxygen consumption was measured at the same growth conditions using a Clark electrode (the data were standardized by cell density). The data indicated that all three strains respired, but at different rates (Figure 1A). The respiration of SEY6210 and W303 was comparable whereas BY4741 exhibited approximately half the rate of the oxygen consumption, a reduction in respiration that could not be explained by growth rates (Figure 1B). BY4741 is known to have a mutation in the gene encoding the Hap1 transcription factor, which regulates the expression of electron transport chain components, thus explaining the observed reduced respiration rate (Gaisne, Becam et al. 1999). As a control, we determined the oxygen consumption rate of a SEY6210 strain deleted for *ATP12*, a gene encoding a chaperone essential for the assembly of the mitochondrial F_O_F_1_ ATP-synthase. This mutant strain exhibited an 80% reduction in respiration, indicating the majority of oxygen consumption by the wild-type yeast strains was linked to mitochondrial ATP production (Figure 1A).

**Figure 1.**
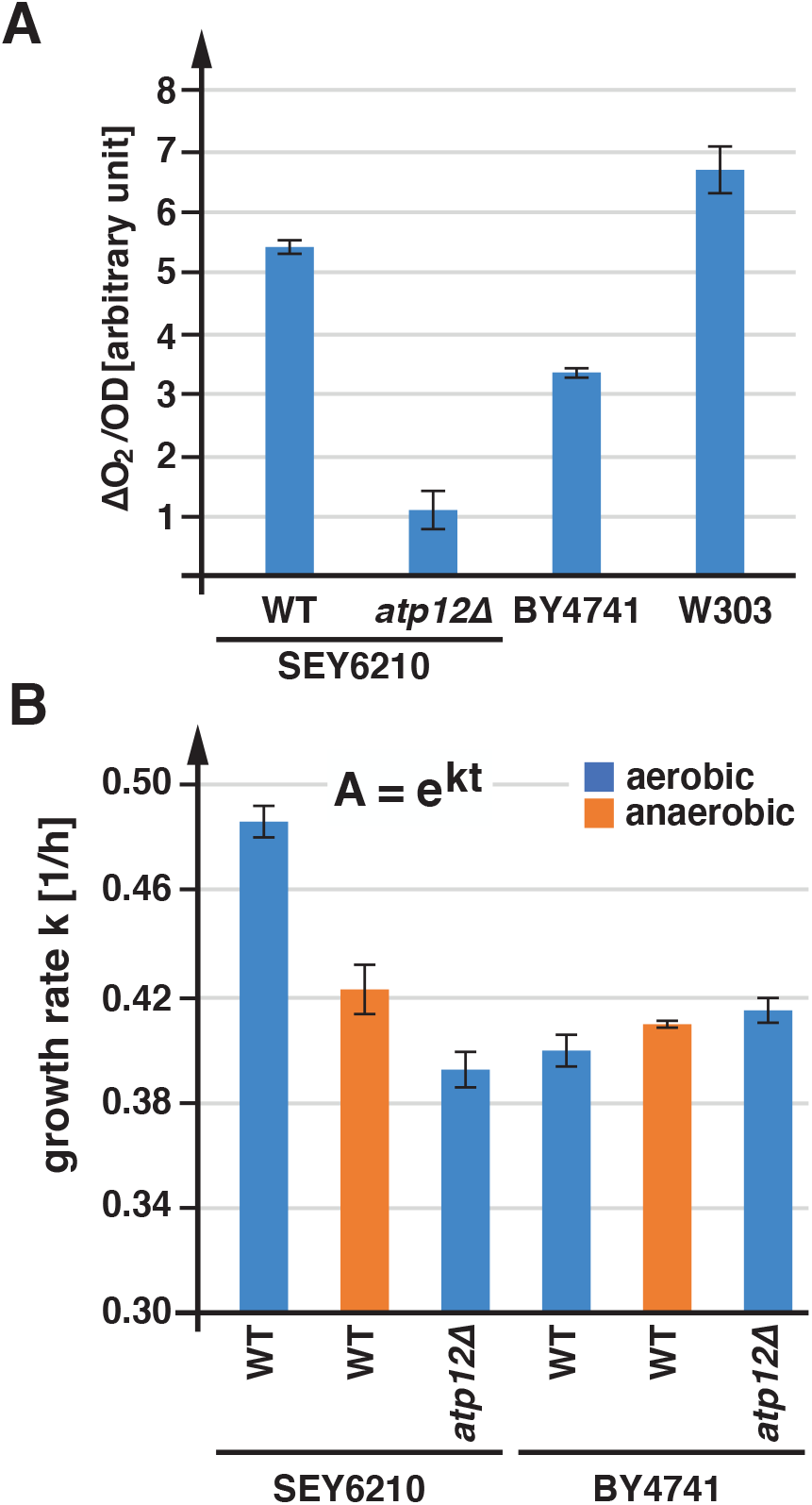
Oxygen consumption and growth rates of lab strains. The bar graphs show the average and standard deviation of 3 measurements. A) Wild-type and *atp12Δ* (AMY36) strains were grown in YPD to and oxygen consumption was measured using a Clark electrode. The data were standardized by the cell density (OD). B) Growth rates in YPD of wild-type and *atp12*Δ (AMY36, YJL180C), in presence or absence of oxygen.

The data in Figure 1A suggested that even at high glucose concentrations, lab yeast strains consume oxygen. To test if this oxygen consumption is important for growth, we compared the growth rates of SEY6210 and BY4741 under aerobic and anaerobic conditions. For SEY6210 the presence of oxygen resulted in a slight growth advantage, whereas BY4741 exhibited better growth anaerobically (Figure 1B). The aerobic growth problems of BY4741 are likely a consequence of the *hap1* mutation. Together, the growth analysis suggested that at high glucose concentrations, respiration contributed a small amount to the growth of yeast, which seemed to support the notion that under these conditions yeast produced ATP mainly by fermentation. However, the *atp12* mutant strain (lacking F_O_F_1_ ATP-synthase activity) of SEY6210 showed a clear growth defect, suggesting that, under aerobic conditions, fermentation was not able to compensate for the lack of respiration. Furthermore, the data indicated that anaerobic growth was improved by metabolic adaptations that supported fermentation.

In the next set of experiments, we determined the oxygen consumption of SEY6120 as a function of the glucose concentration (synthetic complete, SD_com_ medium). As expected, the data showed a strong increase in oxygen consumption at low glucose concentration (0.2%) compared to high glucose conditions (>2%; ~3-fold induction; Figure 2A). It should be noted that at the time the cultures were used for the measurements some of the glucose was already consumed and thus the effective glucose concentration of the samples was at least 0.1% lower than indicated (based on the growth curve of yeast in presence of 0.2% glucose). The de-repression of the oxidative phosphorylation (diauxic shift) is predicted to occur at glucose concentrations below 0.2% (Otterstedt, Larsson et al. 2004). However, our data did not fit to a sharp transition point but indicated a much more gradual adjustment of respiration rates to glucose concentrations. Furthermore, above ~2% glucose, oxygen consumption plateaus at a minimal activity of ~30% relative to the low glucose sample and this activity does not further diminish at higher glucose levels (Figure 2A). This result suggested that the glucose metabolism of SEY6210 is saturated above 2% glucose and never shows complete suppression of respiration.

**Figure 2.**
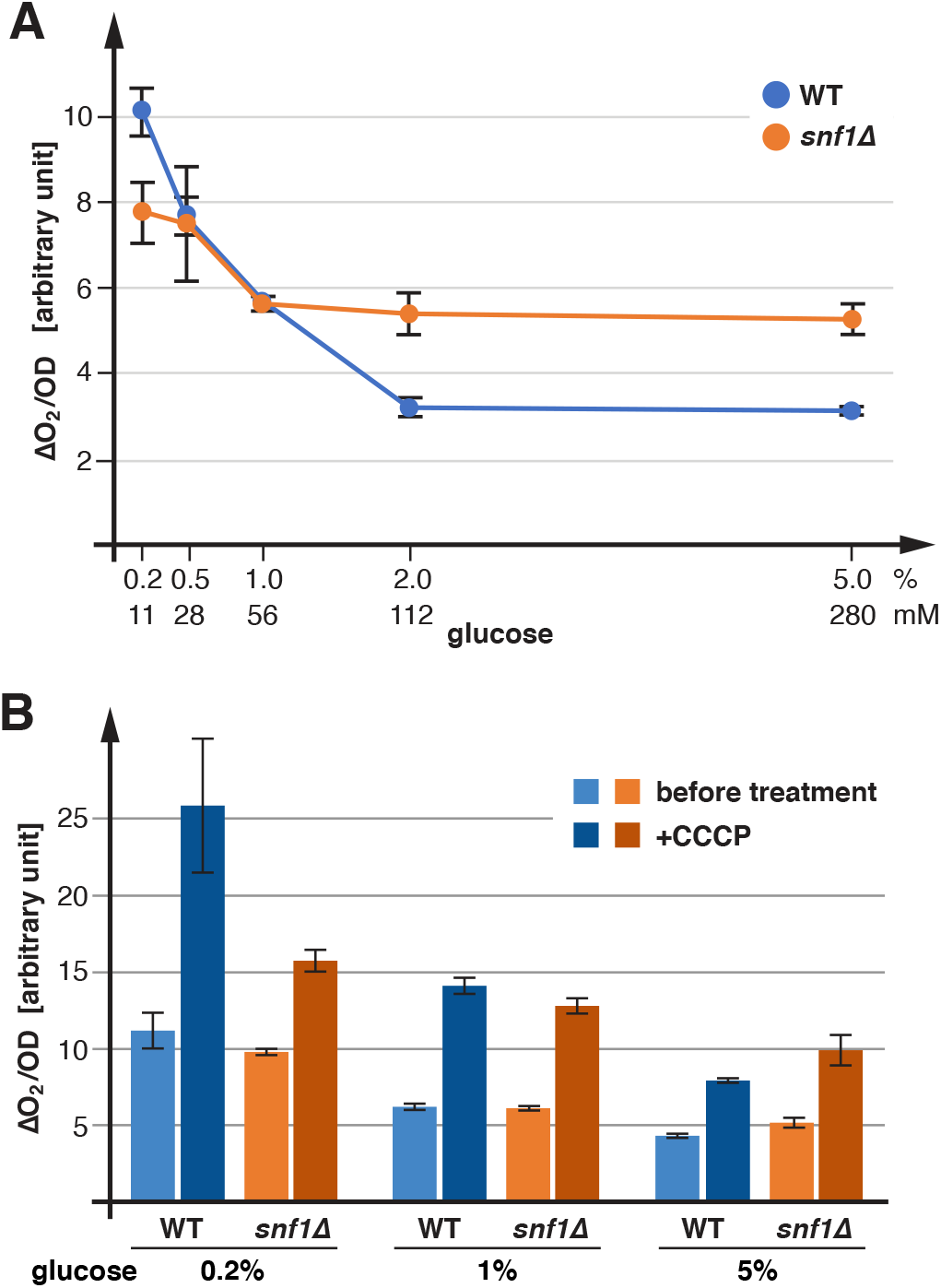
Glucose-dependent oxygen consumption rates of wild type and *snf1Δ* strains. The bar graphs show the average and standard deviation of 3 measurements. A) SEY6210 and AMY20 were grown in SD_com_ containing the indicated glucose concentrations (concentration of the starting growth medium) and oxygen consumption was determined. B) Oxygen consumption of yeast strains (SEY6210, AMY20) grown in SD_com_ in presence of various glucose concentrations. After measuring the initial oxygen consumption rate (before treatment) 50μM CCCP was injected and the resulting increase in oxygen consumption was determined. This data set was obtained independently of the measurements shown in A.

### Respiratory chain substrates are not limiting

It has been proposed that the glucose-dependent changes in oxygen consumption is likely the result of transcriptional regulation of the genes involved in respiration (Kayikci and Nielsen 2015). Alternatively, the competition with fermentation for pyruvate might limit respiration. To test if respiration is substrate limited, we compared oxygen consumption in presence or absence of the proton ionophore carbonyl cyanide *m*-chlorophenyl hydrazine (CCCP), which disrupts proton gradients and thus uncouples oxidative phosphorylation. The results, shown for all glucose concentrations, measured a ~2-fold increase in respiration after addition of CCCP, indicating that the respiratory potential of mitochondria is not substrate limited (Figure 2B). Therefore, the likely culprits limiting oxygen consumption are either the expression levels of F_O_F_1_ ATP-synthase or low levels of ADP. Because a strong proton gradient across the inner mitochondrial membrane is important for protein import into the matrix, the electron transport chain cannot that maintains this proton gradient is unlikely to be limiting.

### Fermentation rates of lab strains

After demonstrating that our lab yeast strains may be respiring more than previously appreciated, we were interested in determining ethanol production as a readout for the energy production by fermentation. We measured by mass spectrometry the ethanol concentration of SEY6210 cultures grown to mid-log phase (OD_600nm_=1.0) in presence of different glucose concentrations. The data indicated that fermentation rates saturate at glucose concentrations >0.5% and only the 0.2% sample showed a substantial drop in ethanol production (~half of maximal; Figure 3A). Compared to the oxygen consumption rates, the rates of ethanol production showed an inverse correlation to the glucose concentration, resulting in a saturation curve of the fermentation/respiration ratio that plateaued at >2% glucose (Figure 3B).

**Figure 3.**
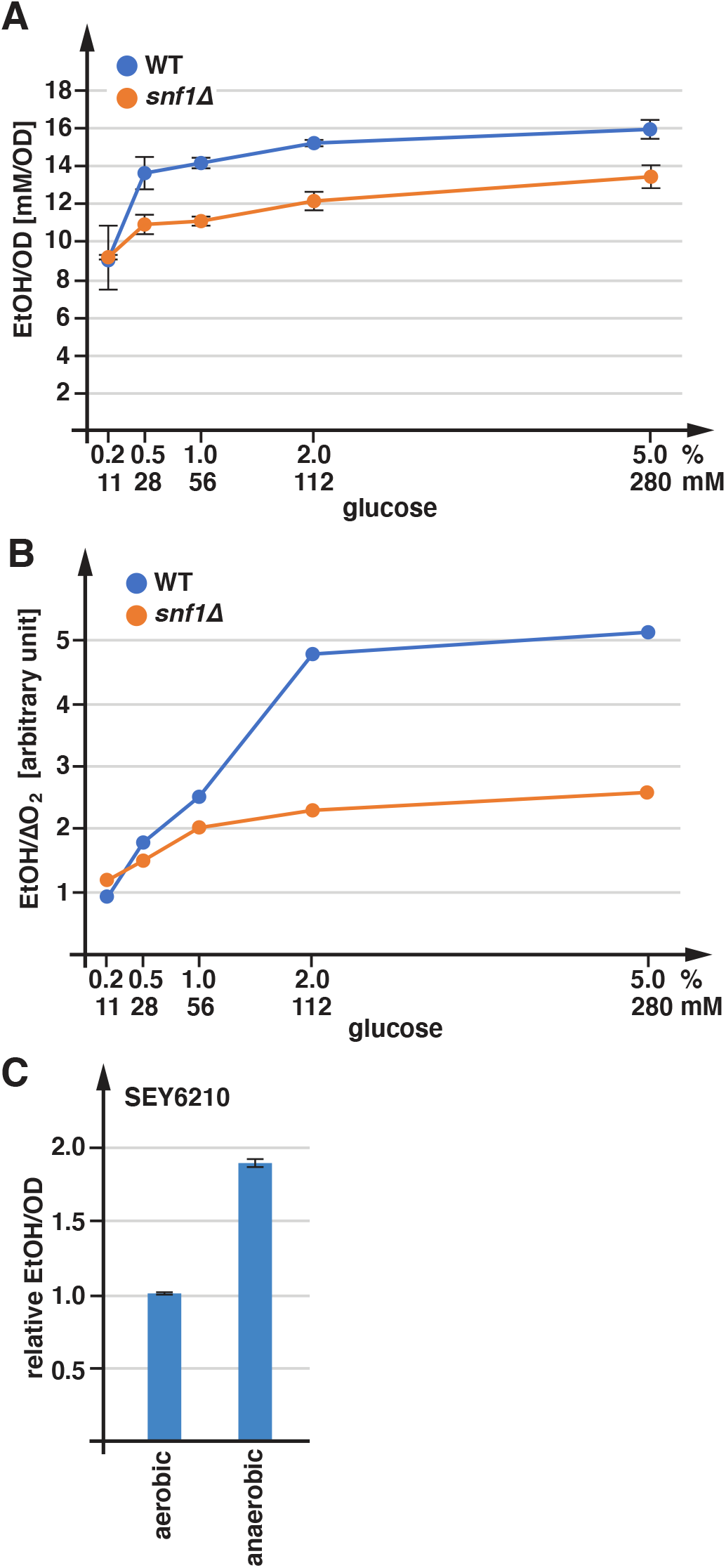
Fermentation rates of wild type and *snf1Δ* yeast strains. The graphs show the average and standard deviation of 3 measurements. A) Yeast strains (SEY6210, AMY20) were grown in SD_com_ in presence of various glucose concentrations. The ethanol concentrations present in the growth medium was quantified by GC-MS and standardized by the cell density. B) Ratio of ethanol concentration to oxygen consumption, using the data sets from Figure 2A and Figure 3A. C) Ethanol production of SEY6210 grown in SD_com_ in presence or absence of oxygen. The data were standardized by the cell density and presented relative to the ethanol production of the aerobically grown cells.

To estimate the contribution of respiration to the energy production of yeast, we compared the ethanol levels of cultures grown anaerobically relative to that of aerobic cultures. We observed that in presence of 2% glucose, ethanol production doubled in anaerobic conditions (Figure 3C). Since anaerobic growth rates are only slightly lower than growth rates under aerobic conditions (Figure 1B), the data indicated that even at glucose-saturation, aerobic cultures produce close to 50% of their energy by respiration. At lower glucose concentrations, the contribution of respiration to the energy production of SEY6210 is even higher (Figure 3B).

### Suppressors of F_O_F_1_ ATPase mutants

We observed reduced growth rates for SEY6210 *atp12*Δ (loss of F_O_F_1_ ATPase), indicating that in this strain glycolysis is not able to compensate for the loss of respiration under aerobic conditions (Figure 1B). Surprisingly, we found that the BY4741 *ATP12* deletion strain we obtained from the knockout collection exhibited no growth defect, but in fact grew even slightly faster than the wild type strain (Figure 1B). We suspected that this strain, which had been propagated as part of the knockout collection for many generations, might have acquired additional mutations that suppress the growth defect caused by the *ATP12* deletion. Therefore, we sequenced the genomes of 4 colonies of the *atp12*Δ strain. In comparison to control sequences, the *atp12*Δ genomes contained several mutations, in addition to the gene deletion that were found in all 4 isolates. Table 1 summarizes the identified mutations that change protein sequences and are of high sequencing quality (supported by >50 reads). Interestingly, we identified a mutation that resulted in a frameshift in the *MIG1* gene. Mig1 is a transcription factor involved in the glucose repression of many metabolic genes, including genes encoding glucose transporters. The *MIG1* frameshift likely represents a loss of function mutation, which is predicted to increase glucose import and glycolysis (Ozcan and Johnston 1995, O’Donnell, McCartney et al. 2015), thereby compensating for the loss of mitochondrial ATP production. Additional mutations in *atp12*Δ were identified in *SLA2* (*END4*), a gene involved in endocytosis (Raths, Rohrer et al. 1993) and in *SEG2*, which encodes an eisosome component. Both of these mutations are predicted to affect the stability of APC-type nutrient transporters, an interesting finding that will be further discussed below. Together, the genomic analysis of *atp12*Δ indicated that mitochondrial ATP production is important for growth of the lab strain BY4741 and that the growth defect of *atp12*Δ is sufficient to give rise to suppressor mutations in a major energy regulatory system.

**Table 1.** Genome analysis of *atp12*Δ mutants from the knockout collection.

|  |  |  |
| --- | --- | --- |
| *MIG1* | Glucose repression | Gaa/Taa: Glu165Stop |
| *SEG2* | Eisosome component | 48bp deletion: Asp599-Asp614 |
| *SLA2* | Endocytosis | tCc/tTc: Ser512Phe |
| *KRE28* | Kinetochore-MT binding complex | Ata/Gta: Ile211Val |
| YPR202W | Putative helicase | Cat/Aat: His7Asn |
| YBL109W | Unknown function | tCt/tGt: Ser65Cys |

### Regulation of respiration and fermentation by AMPK

The finding that deletion of Mig1 is involved in the suppression of the *atp12*Δ phenotype prompted us to determine the role of the Mig1 regulator AMP-dependent kinase (AMPK) in the energy metabolism of our lab strain. AMPK is a highly conserved sensor of ATP homeostasis in eukaryotic cells. In yeast, AMPK has been implicated in glucose repression and therefore in the regulation of fermentation and respiration rates (Kayikci and Nielsen 2015). Yeast AMPK is composed of the kinase Snf1 (α-subunit), the activator Snf4 (γ-subunit) and one of three different ß-subunits (Sip1, Sip2 or Gal83) that localize AMPK to different sites in the cell (Ghillebert, Swinnen et al. 2011). A major downstream target of AMPK is the transcription factor Mig1. Under high glucose/energy conditions AMPK is mainly inactive and Mig1 suppresses the expression of many genes involved in glucose import and the catabolism of alternate carbon sources (Kayikci and Nielsen 2015). Low cellular energy levels cause AMPK phosphorylation and activation, which is mediated by upstream kinases and stabilized by the presence of ADP. Active AMPK phosphorylates and thus inactivates Mig1 and the resulting transcriptional changes support energy production.

To test how AMPK-dependent transcriptional control is involved in regulating the fermentation/respiration ratio, we determined changes in the energy metabolism of the AMPK mutant *snf1*Δ. The *snf1*Δ strain exhibited a drop in oxygen consumption at low glucose concentrations relative to wild type and a 1.7 fold reduction of maximal respiration rate (Figure 2A). At higher glucose concentrations, *snf1*Δ showed increased respiration and reduced ethanol production, which resulted in a lower fermentation-to-respiration ratio (Figures 2A, 3A, 3B). These results indicated that AMPK functions in both the de-repression of respiration at low glucose concentrations and the promotion of glycolysis/fermentation under high-glucose conditions. However, since *snf1*Δ still exhibited regulation of respiration and fermentation (Figure 3B), we conclude that AMPK is not the key sensor in this glucose-dependent regulation, rather it acts as an enhancer of this adaptive response. This effect of AMPK on the fermentation/respiration ratio might be explained by the fact that deletion of *SNF1* lowers the levels of the low-affinity glucose transporters Hxt1 and Hxt3 (O’Donnell, McCartney et al. 2015), which reduces glucose flux and shifts the import function to the high-affinity glucose permeases Hxt2/4. Consequently, the glucose saturation concentration shifted from 2% in wild type to ~1% in *snf1*Δ (Figure 2A). The intracellular glucose sensor that provides respiration regulation in *snf1*Δ might be Hxk2, a hexokinase that also functions in the regulation of Mig1 targets (Vega, Riera et al. 2016).

### Energy levels affect nutrient import

The energy levels are expected to first and foremost affect cellular systems that are energy intensive, such as the APC transporter–Pma1 system. Pma1 is the ATP-driven proton pump that is responsible for maintaining a strong proton gradient across the plasma membrane. With over 10^6^ molecules per cell, Pma1 is the most abundant protein in yeast and is predicted to represent one of the main ATP consumers (Perlin, Harris et al. 1989, Cyert and Philpott 2013). It is therefore not surprising that Pma1 activity is tightly controlled by the metabolic state of the cell (Goossens, de La Fuente et al. 2000). Because of its function in pH and ion homeostasis, Pma1 is an essential protein. The 26 different APC-type transporters in yeast are major consumers of the proton gradient, which use this gradient to import small molecule nutrients such as amino acids and nucleobases. Changes in ATP levels are expected to affect the strength of the proton gradient, which in turn affects nutrient import. Therefore, we measured the import activity of an APC transporter as a readout for the cellular energy levels.

The kinetics of uracil import by the APC-transporter Fur4 is dependent on the uracil concentration in the growth medium, the number of Fur4 transporters and the strength of proton gradient. Therefore, by keeping the concentration of uracil and Fur4 constant we were able to measure the strength of the plasma membrane proton gradient in different yeast strains and growth conditions. For these experiments we grew the yeast strains in uracil-free medium to mid-exponential growth phase, added 20μg/ml uracil for 10min and after washing with ice-cold water the cells were extracted with methanol at 50°C which was injected into the HPLC system for analysis. We measured uracil import in strains expressing a plasmid-encoded Fur4(ΔN)-GFP, an N-terminally truncated form of Fur4 that is not ubiquitinated and thus is expected to maintain similar expression levels in the different strains. In addition, we quantified the expression levels of Fur4(ΔN)-GFP by flow cytometry and used the average fluorescence intensity to standardize the uracil import data. It should be noted that the strains also express the genomically encoded *FUR4* gene and thus the measured import activity is the sum of both transporters, Fur4 and Fur4(ΔN)-GFP. However, SEY6210 without the *fur4(*Δ*N)-GFP* plasmid exhibited ~10-fold lower uracil import activity (data not shown) and therefore we ignored the contribution of the genomic encoded Fur4.

We found that in SEY6210 expressing Fur4(ΔN)-GFP, inhibiting ATP production by the addition of NaN_3_/NaF (blocks ATP production by oxidative phosphorylation and glycolysis) 5 min before the addition of uracil, almost completely blocked uracil uptake (Figure 4A), suggesting that the proton gradient across the plasma membrane collapses rapidly after losing ATP and thus Pma1 activity. Blocking mitochondrial ATP synthesis by deleting *ATP12* resulted in a ~50% drop in uracil uptake (Figure 4A), indicating that oxidative phosphorylation is important to maintain a strong proton gradient. In contrast, loss of AMPK activity (*snf1*Δ) did not seem to affect the proton gradient, suggesting that this strain is not energy starved but, based on the data in Figure 3B, shifted part of ATP production from fermentation to respiration. BY4741 strains exhibited generally lower uracil uptake, consistent with the respiration defect observed in this strain (see Figure 1A). BY4741 *atp12*Δ showed increased uracil import compared to the parental wild-type strain, indicating that the acquired suppressor mutations indeed compensated for the loss of mitochondrial ATP production (Figure 4A). Finally, anaerobic growth of SEY6210 reduced uracil import activity to a similar extent as deleting *ATP12* (Figure 4B), suggesting that anaerobically growing cells do not fully compensate for the lack of respiration and thus maintain a lower proton gradient. However, in rich media, this reduced proton gradient had no major effect on growth (Figure 1B).

**Figure 4.**
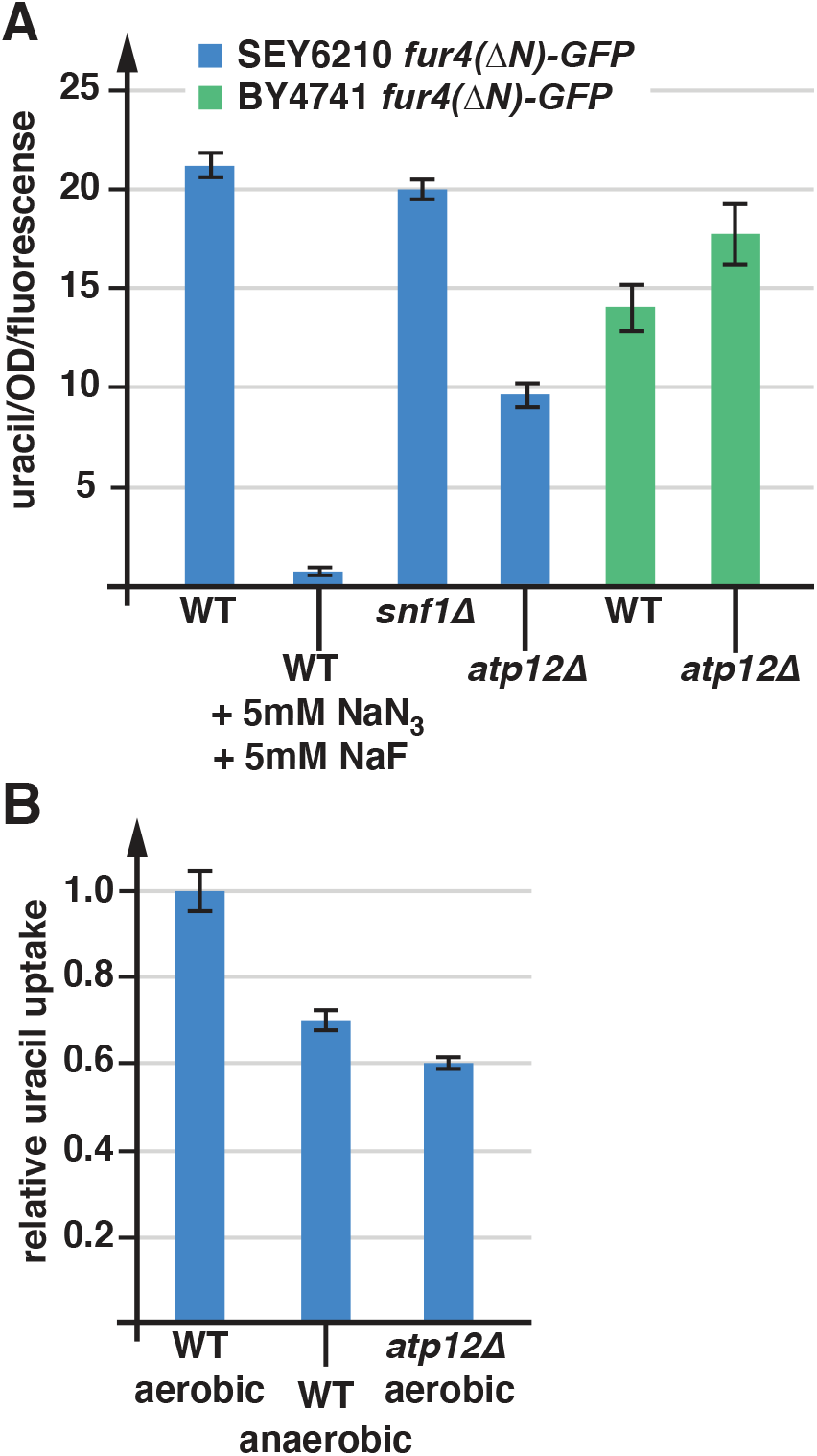
Uracil import is affected by the energy state of the cell. The bar graphs show the average and standard deviation of 3 measurements. A) For the uracil uptake assays the strains were grown in SD_com-ura_ medium. The following strains were used for these experiments: WT (in blue; SEY6210 pJK88), *snf1*Δ (in blue; AMY20 pJK88), *atp12*Δ (in blue; AMY36 pJK88), WT (in green; BY4741 pJK88), *atp12*Δ (in green; YJL180C pJK88). B) Relative uracil uptake of wild-type and *atp12*Δ (SEY6210 pJK88, AMY36 pJK88) grown under aerobic or anaerobic conditions.

### Response to an acute drop in respiration

The data discussed above suggested that fermentation is not able to compensate for the lack of respiration. Therefore, we would expect that the acute block of respiration will cause a dramatic drop in ATP levels that cannot easily be compensated for by increased fermentation. If energy production cannot be increased, the cells might respond to the low ATP levels with changes on the energy demand side. Consistent with this idea, after addition of NaN_3_ (blocks oxidative phosphorylation) or oxygen depletion we observed an increase in the turnover of the APC transporters Fur4 and Mup1 (GFP signal in the vacuole and small intracellular structures, most likely endosomes; Figure 5A). This finding suggested that yeast responded to the loss of respiration by reducing APC transporter activity, which lowers the proton-flux and consequently lowers the energy demand of Pma1. Other transport systems that are not proton driven, such as the glucose transporter Hxt3 or the iron transporter Ftr1, did only weakly respond to the energy loss (Figure 5A). Together, the data suggested that in comparison with other nutrient importers, APC transporters respond particularly sensitively to a drop in energy levels. However, unlike an acute change in energy production (Figure 5A), cells adapted to the conditions did not show an increase in Fur4 downregulation (Figure 5B), suggesting that APC transporter downregulation occurs during the adaptation period and does not persist.

**Figure 5.**
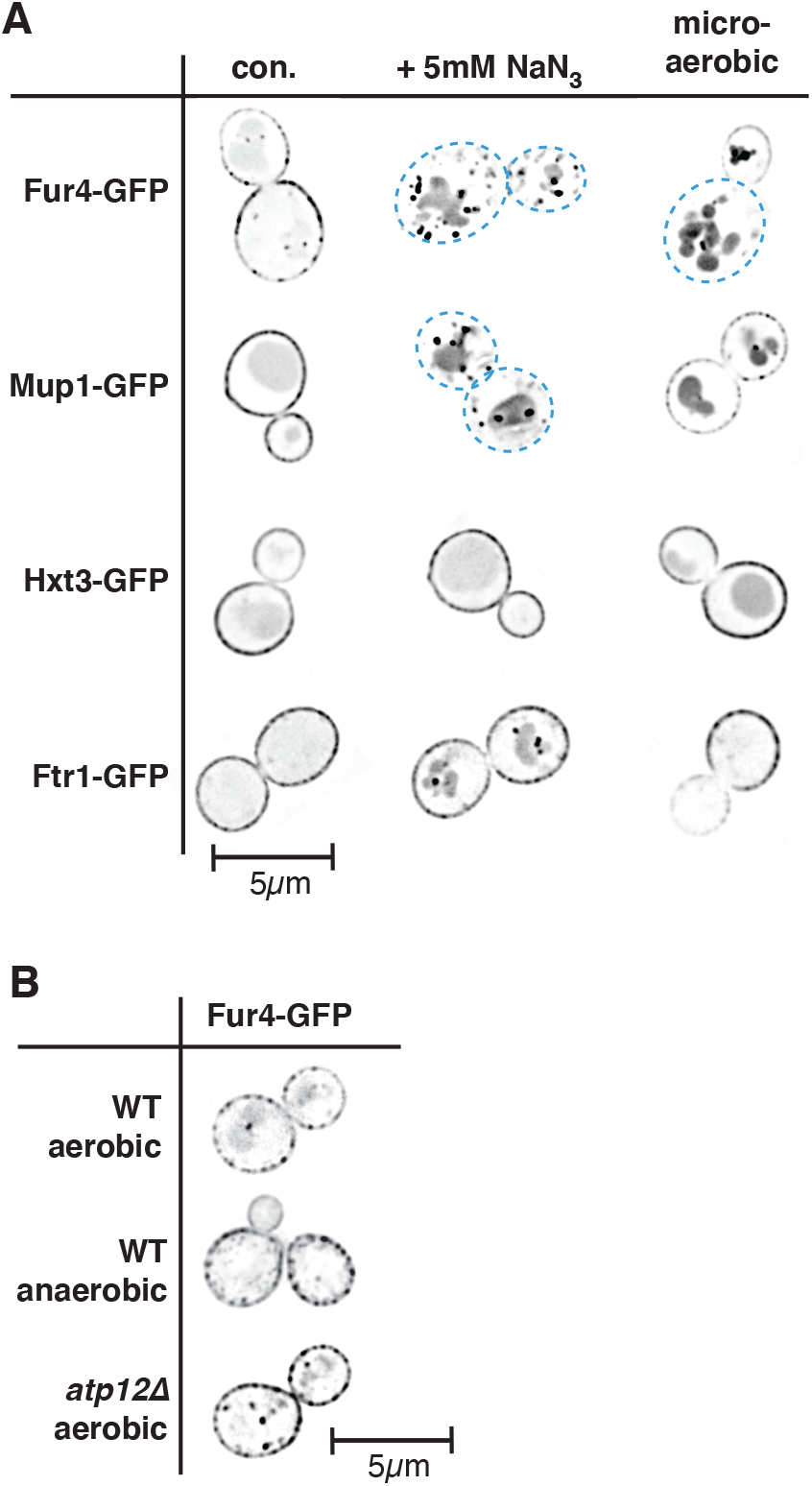
ATP levels affect the localization of APC transporters. A) Yeast strains were grown in SD_com_, SD_com-ura_ or SD_com-leu-met_ medium to mid-log phase, treated for 1h as indicated and analyzed by fluorescence microscopy (con.: control before treatment). The pictures show one optical section from the middle of the cell. The pictures were inverted for better visualization of the GFP signal (GFP signal is shown in black). Blue dashed line indicates the cell outline where surface expression of the GFP-tagged protein is low. The following strains were used: Fur4-GFP (SEY6210 pJK19), Mup1-GFP (SEY6210 pPL4146), Hxt3-GFP (DLY050), Ftr1-GFP (CBY118). B) Fluorescence microscopy of yeast strains grown in SD_com-ura_ expressing Fur4-GFP (single cross section). The imaged strains are: WT (SEY6210 pJK19), *atp12*Δ (AMY36 pJK19).

## Discussion

We originally started this study with the goal to better understand the downregulation of APC transporters observed during different stress conditions. A key finding from our experiments is that during many stress conditions APC transporters might be downregulated because of stress induced changes in energy metabolism. A likely reason for this regulation is the high ATP demand of the plasma membrane proton pump Pma1, which maintains the proton gradient that drives nutrient import by the APC transporters. Balancing proton import and export is essential to prevent acidification of the cytoplasm. A drop in ATP levels reduces proton export by Pma1 and as a result cells reduce proton import by downregulating APC transporters.

One condition that causes APC transporter degradation is the switch from well aerobic to micro-aerobic growth, which suggested that respiration is important to maintain Pma1 activity. This observation was unexpected since at the high glucose concentrations used in our growth medium (2% glucose), *S. cerevisiae* has been shown to rely mainly on glycolysis/fermentation for ATP production (e.g. (Fendt and Sauer 2010)). This aerobic fermentation is also referred to as the “Crabtree effect” (a variation on the Warburg effect), which describes an energy metabolism that even in presence of oxygen shunts most of the imported glucose to ethanol production and suppresses respiration. However, strains used for metabolic studies often differ significantly from the strains used in cell biology and even among those strains, genetic diversity has complicated the view of yeast metabolism (as shown for the respiratory defective strain BY4741; Figure 1A). We found that SEY6210 (and W303), the yeast strain we commonly used for our trafficking studies, exhibited robust respiration with rates that did not drop below 30% of maximum even at high glucose concentrations. This result suggested that even at high glucose levels ~1/3 of the ATP production was generated by respiration (assuming that at glucose concentrations <0.2% most of the ATP is generated from respiration). Compared to aerobic conditions, ethanol production doubled under anaerobic, high glucose growth. This result suggested that respiration might contribute half of ATP production in standard lab growth conditions. The discrepancy between these two numbers (1/3 versus half of ATP production) might be explained by the observations that under anaerobic conditions yeast grew slower and possessed a weaker proton gradient (Figures 1B and 4B), indicating that these cells exhibited lower ATP production rates. Therefore, 50% of anaerobic ATP production might be equivalent to ~30% of ATP production rate under aerobic conditions.

Together, our data demonstrate the importance of respiration for the energy metabolism of laboratory yeast strains (Figure 6). Disrupting oxidative phosphorylation by removing oxygen or deleting the ATP synthase assembly factor Atp12 resulted in reduced growth and a weaker proton gradient. The impact of the *ATP12* deletion on growth was severe enough to generate a selection pressure for suppressor mutations, which included a loss of function mutation in *MIG1*, a mutation that increases glucose uptake and is able to compensate by increasing fermentation rates (see Table 1) (Kayikci and Nielsen 2015). It is interesting to note that the *mig1*Δ strain grew better and exhibited a stronger proton gradient than the corresponding wild type strain BY4741, suggesting that regulatory mechanisms in BY4741 are not able to compensate for the reduced respiration rate of this strain (BY4741 is mutated in *HAP1* which reduces respiration by 50%; Figure 1A). In general, we were surprised to find that fermentation was not able to fully compensate for the lack of respiration, even after extended adaptation period (several hours of anaerobic growth). Furthermore, yeast was not able to rapidly compensate for an acute loss of respiration, suggesting that under normal growth conditions the fermentation flux is close to its maximal capacity. As a consequence, yeast responds to a drop in ATP by shutting down high energy consumers such as the nutrient transporter/Pma1 system, which is not essential for the cell’s survival (yeast is able to synthesize all amino acids from glucose).

**Figure 6.**
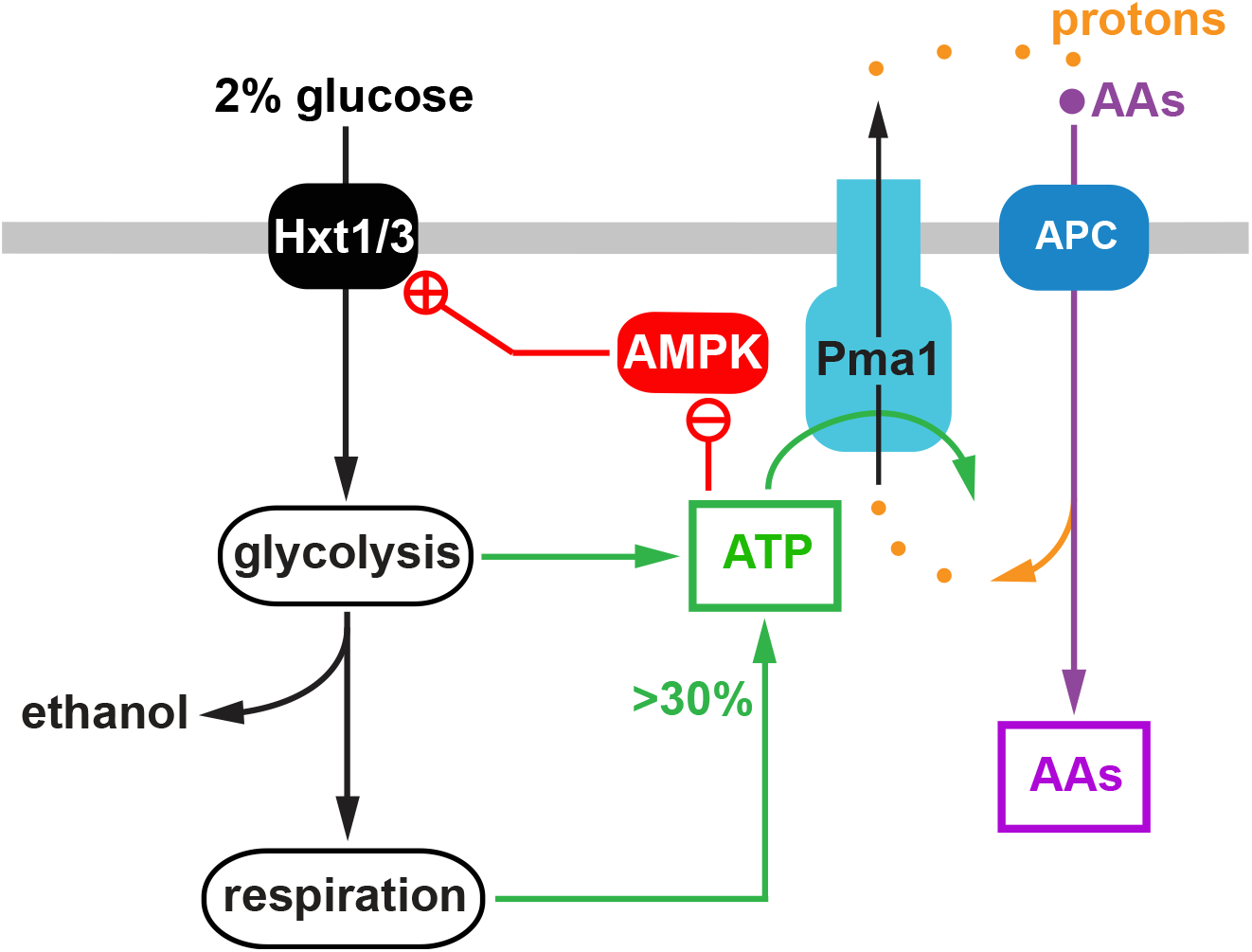
Model of the link between the APC transporter system and the energy metabolis of yeast. The APC transporters require a strong plasma membrane proton gradient for their function (import of amino acids (AAs)), which is maintained by the ATP-dependent proton pump Pma1. ATP required for Pma1 activity is synthesized by respiration (providing 30-50% of ATP) and glycolysis/fermentation. AMPK is regulated by the energy status of the cell and its activity affects the stability of the glucose transporters Hxt1 and Hxt3.

In yeast, AMPK senses energy levels and accordingly phosphorylates downstream targets such as the transcription factor Mig1 (Kayikci and Nielsen 2015). The goal of this regulation is to adjust the energy metabolism to varying glucose levels. We found that a major effect of *SNF1* deletion was reduced fermentation and increased respiration rate at high glucose concentrations, indicating that AMPK is a strong promoter of the Crabtree effect. A likely explanation for this activity of AMPK is the fact that this kinase regulates the stability of glucose transporters at the plasma membrane (O’Donnell and Schmidt 2019) (Figure 6). AMPK phosphorylates and inhibits a subset of alpha-arrestins, adaptor proteins that bind to glucose transporters and recruit the ubiquitin ligase Rsp5. Rsp5 ubiquitinates the transporters which results in endocytosis and degradation of these proteins. Therefore, high AMPK activity is able to stabilize glucose transporters at the cell surface. Loss of AMPK (*snf1*Δ) reduces the numbers of glucose transporters at the cell surface, thereby reducing glucose flux and thus suppressing the Crabtree effect. Consistent with this observation, a previous study showed that reducing glucose import is able to convert a Crabtree-positive yeast into a Crabtree-negative strain (Otterstedt, Larsson et al. 2004).

In summary, this study highlights the importance of oxidative phosphorylation for the energy metabolism of yeast strains commonly used in cell biological studies. Because the energy metabolism is linked to many cellular processes, including the regulation of nutrient transporters, it is essential to consider the growth conditions when designing cell biological experiments. In particular, changes to glucose concentrations or aeration of the yeast cultures should be avoided.

## Materials and Methods

*S. cerevisiae* strains and plasmids used in this study are listed in Table 2. Genomic deletion and tagging were made by homologous recombination as described previously (Longtine, McKenzie et al. 1998). The resulting insertions or gene replacements were confirmed by PCR. The yeast strains were grown either in rich YPD (yeast extract, peptone, and 2% dextrose) or Synthetic Dextrose (SD) medium (yeast nitrogen base; 2% glucose, unless stated otherwise) either with amino acids (SD_com_, in mg/L: p-Aminobenzoic acid (7.5), Ala (75), Arg (75), Asp (75), Asn (75), Cys (75), Glu (75), Gln (75), His (75), Ile (75), Leu (376), Lys (75), Met (75), Phe (75), Pro (75), Ser (75), Thr (75), Trp (75), Tyr (75), Val (75), Adenine (20), Inositol (75), and Uracil (75)). Depending on plasmids present in the strains the SD_com_ medium lacked certain nutrients (-uracil, -leucine). For microscopy experiments with Mup1-GFP expressing strains the SD_com_ medium lacked methionine. To induce expression of constructs containing the *CUP1-1* promoter 0.1 mM cupric sulfate was added to the medium.

**Table 2.**
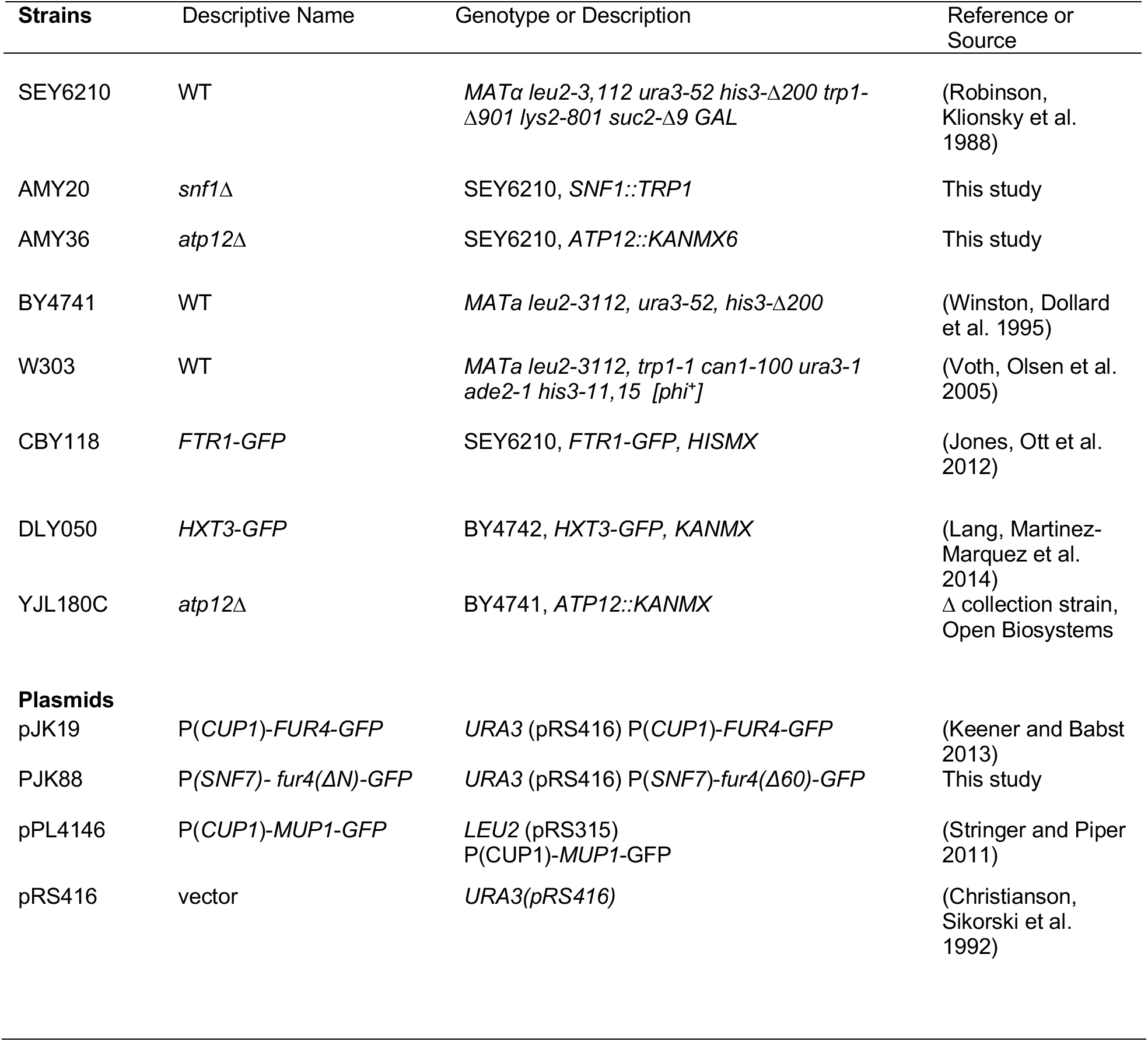
Strains and plasmids used in this study

### Fluorescence microscopy

For microscopy, cells were grown overnight and then diluted into fresh medium in the morning to OD_600_=0.1. The cells were analyzed in exponential growth phase (OD_600_=0.6-0.8). For the microscopy a deconvolution microscope (DeltaVision, GE, Fairfield, CT, USA) was used.

### Uracil uptake assay

For the uracil uptake assay, cells were grown in SD_com-ura_ media to mid log phase and 20μg/mL uracil was added to the cells for 10 minutes after which cells were kept on ice for 5 minutes. Cells were washed three times with ice cold water and resuspended in 50 μl of methanol. They were heated to 55°C for 10min, spun down and the supernatant was separated on an HPLC system using a Luna-NH2 column (Phenomenex) in presence of a 100%-80% acetonitrile/water gradient. The presence of uracil was detected by absorption spectroscopy at 260nm.

### Oxygen consumption

Cells were grown to mid-log phase and oxygen consumption was measured at 30°C with a Clark electrode (Ocean Optics FOXY-R) and data was collected using the OOI sensors software program. Data were normalized according to OD_600_ of the individual cultures.

### Determining ethanol production

Mid-log phase yeast cells grown in SD complete media at 2% glucose were centrifuged at 14,000 x rpm for 2 minutes, and media fraction was collected, and kept in a sealed tube on ice. Just prior to run on GCMS, 0.05% C13-ethanol (Sigma) was spiked into the collected media. Analysis of C12:C13 ratio was done by comparing masses of 45.1 for C12, and 47.1 for C13. The samples were analyzed with a Thermo Scientific TRACE 1310 gas chromatograph coupled to a Thermo Scientific ISO 7000 mass spectrometer equipped with an EI ion source. A 30 m x 0.25 mm I.D. x 0.25 μm Thermo Scientific TraceGold TG-XLBMS column was used to retain EtOH. The GC was set in splitless mode. The helium carrier gas flow rate is 0.8 mL/min. The oven temperature gradient is: 0-2 min; 40°C, 2 - 6.2 min; 120 ℃ with 25°C/min rate. Ion source temperature is 230°C. The mass scan range is between 45 m/z to 60 m/z.

### Genome sequencing

Genomic DNA purification was performed from 4 single colonies of the *atp12*Δ strain from the yeast knockout collection (BY4741 collection, Open Biosystems) using the Qiagen DNeasy Blood and Tissue Kit. Sequencing was performed by Novogene using Illumina HiSeq platform (150 bp paired end sequencing). The resulting data were aligned to the S288C reference genome using BWA-MEM. Variants were called using the GATK tool HaplotypeCaller. Both BWA-MEM and HaplotypeCaller were run with default parameters. The effects of variants were determined using SnpEff and the gene annotation version R64-1-1.81. Downstream filtering of variants to find unique and common ones among groups was performed with unpublished MATLAB and R scripts.

